# A Human Precision Cut Kidney Slice Platform for Studying Glomerular Biology and Therapeutic Responses

**DOI:** 10.64898/2026.10.16.759734

**Authors:** Shreeram Akilesh, Xianwu Li, Bre Mills, Chetan Poudel, Pedro Vasquez, Emily Beirne, Joshua C. Vaughan, Kelly D. Smith

## Abstract

**Introduction:** Short term human precision cut kidney slice cultures (PCKS) represent a tractable experimental that contains mature kidney cell types within their native anatomic context and that can be manipulated *in vitro*. In this study, we employ genomic tools to analyze the PCKS response to culture conditions and their suitability for study of glomerular disease biology.

**Methods:** The impact of culture medium and oxygenation on PCKS viability and gene expression was explored using spatial transcriptomics and traditional histologic approaches. The biological response of PCKS to IFNγ stimulation and signaling suppression by the JAK1/2 inhibitor baricitinib were also explored.

**Results:** A genome wide atlas of glomerular and tubulointerstitial gene expression patterns was generated from PCKS cultured in different media and oxygenation conditions and at various timepoints in culture. IFNγ induced APOL1 gene and protein expression in glomeruli, which could be suppressed by baricitinib. Expression changes were reproducible in PCKS derived from multiple donors.

**Conclusions:** PCKS are reproducibly responsive to IFNγ stimulation, which will allow dissection of APOL1 mediated glomerular disease mechanisms in a human tissue model system. Spatial transcriptomic profiling of tissue responses helps to tailor culture medium/oxygenation to optimize the PCKS system to address specific biological questions.

**Translational Statement:** Human kidney slice cultures preserve mature cells within their native tissue architecture, enabling direct study of disease mechanisms and drug responses in human tissue. Combining this platform with spatial transcriptomics identified reproducible tissue responses to the inflammatory cytokine IFNγ. It also uncovered expected and unanticipated effects of baricitinib, a clinically approved medication. This kidney slice approach could help prioritize therapies for glomerular diseases, including APOL1-associated kidney disease, and identify potential adverse effects before patient trials. This platform offers a fully human tissue system for preclinical testing of candidate therapies and assessment of their molecular effects within the native kidney microenvironment.

## INTRODUCTION

Much has been learned about the anatomy and physiology of the kidney in animal model systems. These experimental models allow for careful dissection of timing and mechanism of kidney disease processes. However, animal models may not recapitulate all aspects of human disease due to differences in physiology and genetics. Recent notices from the National Institutes of Health and the Food and Drug Administration’s (FDA) Modernization Act 2.0, recognize this limitation and urge the use of physiologically relevant human *in vitro* models as alternatives to animal-based experimentation^1, 2^. Kidney organoid models and microphysiological systems (aka “kidney-on-a-chip”) are tractable human model systems to study human disease processes^3^. However, they too have drawbacks such as cellular immaturity, lack of key cell types (e.g., endothelial cells, innate immune cells) and low architectural complexity.

Precision cut kidney slice cultures (PCKS) represent valuable alternative experimental systems which have also contributed to our understanding of renal physiology. In this approach, vibratome-cut ∼200-500µm thick sections of fresh kidney tissue are cultured *in vitro* for up to 72 hours followed by biochemical, histochemical, histologic or other functional analyses. Kidney tissue is most often sourced from patients undergoing nephrectomy for removal of a renal tumor. Some of the earliest reported experiments studied the differences in metabolism between neonatal and adult kidney utilizing slice cultures from multiple animal species and humans^4–6^.

Studying the response of rabbit cortical kidney slices to dog plasma contributed to the discovery of atrial natriuretic peptide as a hormonal factor that promotes sodium excretion by nephrons^7^. Rat kidney slice cultures were also used to establish the mechanisms of amino acid transport in proximal tubules^8^. The ability to generate replicate slice cultures is a valuable design feature that allows for experimental perturbations and time course studies, which has been leveraged to study cisplatin toxicity, the maladaptive effects of dexamethasone on tubular morphology and function, and the mechanisms of fibrosis^9–19^.

In contrast to above mentioned studies of tubular physiology and function, glomerular focused slice culture studies have primarily used imaging-based analyses. One early study used a calcium reporter dye, Fluo-4, to study calcium signaling in mouse glomeruli using 100µm thick cortical slices^20^. Using kidney slices from genetically modified mice, investigators have used multi-photon microscopy to study calcium signaling in podocytes and super-resolution microscopy to study the ultrastructure of the glomerular slit diaphragm^21, 22^. Combining tissue clearing approaches with the application of biomolecule-reactive fluorescent probes permits high resolution studies of kidney architecture^23, 24^. The recent advent of spatial transcriptomics technologies opens up the possibility of deep, spatially registered transcriptomic profiling of slice cultures. Therefore, in this study, we use PCKS cultures to explore human glomerular disease mechanisms using a combination of conventional histology, lightsheet microscopy, and spatial transcriptomics. We demonstrate that PCKS exhibit good viability in short term (<72 hours) *in vitro* culture and reproducible biologic responses after cytokine stimulation, making them a powerful platform for studying glomerular disease mechanisms.

## METHODS

### Human tissue collection

The work described has been carried out in accordance with The Code of Ethics of the World Medical Association (Declaration of Helsinki) for experiments involving humans. Potential donors undergoing nephrectomy for removal of kidney tumors were identified using the University of Washington’s NWBioSpecimen tissue procurement service. After obtaining informed consent under the University of Washington’s IRB Study Protocol 4416 and Fred Hutch IRB Study Protocol 7397, the nephrectomy specimen was triaged by NWBioSpecimen pathology assistants, who provided deidentified pieces of uninvolved kidney cortex (∼5-10g) to the study team in cold RPMI medium. Ischemia time to sample hand off was ∼1 hour. Patient clinical characteristics and pathologic findings are summarized in **Supplemental Table 1**.

### Kidney precision cut slice cultures

Kidney slice culture was performed using a protocol previously described for liver tissues^25^. Cores of kidney cortex were generated using 6 mm diameter biopsy punches and placed into cold HBSS (Gibco 14025092). After blotting excess buffer with sterile gauze, cores were embedded in low gelling temperature 2% agarose (Sigma A0701) made in complete culture medium (see composition below). The agarose embedded cores were placed on ice and solidified within 10 minutes. Using cyanoacrylate (Ted Pella 16024), the bottom of the embedded core was glued directly to the vibratome specimen holder disc. After loading the specimen disc and HBSS-containing buffer tray into the vibratome (Leica VT 1200S), the space around the buffer tray was filled with ice. Vibratome sections were obtained at a thickness of 250 µm, a cutting speed of 0.8mm/s and amplitude of 2-3. After slices were floated onto the buffer, one slice was placed into each well of a 24-well dish containing 450 µl of prewarmed complete medium, containing a 0.4µm cell culture insert (Millipore PICM0125) and overlaid with additional 150 µl complete medium. On average, it took an additional 2 hours after tissue procurement to prepare slices for culture; the time of initiation of culture was considered the T0 timepoint. Slice cultures were then incubated at 37°C in a 5% CO_2_ atmosphere or 95% O_2_/5% CO_2_ (carbogen atmosphere) for the indicated time periods.

Two media formulations were tested. Medium A consisted of William’s medium E (Gibco A1217601) containing 5% fetal bovine serum (Hyclone SH30087.02), 0.5% glucose (Gibco A2494001), 100 unit/ml penicillin and 100 µg/ml streptomycin (Gibco 15140148) and 15 mM HEPES (Gibco 15630106). Medium B consisted of DMEM/F12 (containing 0.5% Glucose) supplemented with 100 U/ml penicillin, 100 µg/ml streptomycin, 15 mM HEPES, Glutamax, Insulin-Transferrin-Selenium (ITS), 1mM glutathione, 10 U/ml superoxide dismutase, and MEM Non-Essential Amino Acids (NEAA). For selected experiments, kidney slices were treated with 100 ng/ml human IFNγ (R&D Systems 285-IF-100/CF) and/or 1 or 10µM baricitinib (SelleckChem S2851)^26^. Medium alone or an equivalent volume of DMSO was used as controls for baricitinib treatment experiments. For cytokine stimulation or small molecule treatment, the reagents were premixed with the culture media and placed in the tissue culture dish prior to addition of the tissue slice.

### Assessment of slice viability

Cell death was assessed by measuring cytoplasmic lactate dehydrogenase (LDH) release into the culture supernatant using an LDH Cytotoxicity Assay kit (Cell Signaling 37291). Maximal release was determined using replicate slices incubated with 1% TX-100. 100µl of supernatant from each slice culture was incubated in the dark with LDH reaction solution for 30 minutes at 37°C with gentle shaking. Well absorbance at 490 nm was measured using a plate reader after subtracting average background absorbance contributed by blank wells. Percent viability was calculated as (1 - (sample - media background) / (triton X-100 - media background)) x 100.

### Quantitative reverse-transcription PCR (qRT-PCR)

Total RNA was isolated using RNeasy Mini Kit (Qiagen #74104). cDNA synthesis was performed by GoScript Reverse Transcriptase Kit (Promega #A5000). Quantitative PCR was carried out with iTaq Universal SYBR Green Supermix (Bio-Rad #1725121) using QuantStudio 6 Flex Real-Time PCR System (Thermo Fisher Scientific). Expression levels were normalized to GAPDH, and fold change was determined by application of the 2^-ΔΔCT^ method. The sequences of the qPCR primers used are provided in **Supplemental Table 2**.

### RNA in situ hybridization (RNA-ISH) and APOL1 immunohistochemistry

Tissue slices were fixed in 10% neutral buffered formalin for 24 hours and then transferred to 70% ethanol for another 24 hours at 4°C. Samples were then paraffin embedded and *en face* sections were cut at 4 µm thickness and affixed to SuperFrost Plus charged microscope slides. Manual chromogenic RNA-ISH was performed according to the manufacturer’s instructions using an ACDBio RNAscope 2.5 HD RED Reagent Kit (Catalog# 322350) and probes targeting the *APOL1* gene (Catalog# 459791).

For detection of APOL1 protein in tissue sections, slides were deparaffinized, rehydrated, and subjected to a 20-minute incubation of 100°C Tris-based pH 9 antigen retrieval solution (catalog H-3301, Vector Labs, Newark, CA). Following a 20-minute cooling period, slides were blocked with 3% hydrogen peroxide for 10 minutes and incubated in 2.5% normal horse serum block (catalog 30022, Vector Labs, Newark, CA) for 20 minutes. Slides were incubated overnight with rabbit anti-APOL1 antibody (1:5,000 dilution, catalog HPA018885, Millipore Sigma, Burlington MA). Antibody detection was completed via a 30-minute incubation with horse anti-rabbit IgG polymer (ImmPress HRP detection kit, catalog MP-7401, Vector Labs, Newark, CA) and 3-minute DAB development period (ImmPact DAB substrate kit, catalog SK-4105, Vector Labs, Newark, CA). Slides were counterstained with Harris hematoxylin (catalog 6765001, Epredia, Kalamazoo, MI), dehydrated, cleared, and mounted.

### Kidney slice FLARE staining and tissue clearing

Fixed human kidney slices of 250 µm thickness were stained with FLARE stains with slight modifications to the published protocol^23, 24^. Only 2 of the tri-color stains were used. Oxidized carbohydrates were stained with 1 mg/ml concentration of AT565-hydrazide (fluorescent analog of the PAS stain) for 30 minutes at room temperature. Nuclei were labeled with 3.5 mg/mL concentration of SYBR Green I dye for 2 hours in a 50% tetrahydrofuran (THF) and 50% PBS mixture at 4°C. Tissues were dehydrated using a graded series of THF:water mixtures and a short incubation in dichloromethane (DCM) before refractive index matching in ethyl cinnamate.

### Lightsheet microscopy and image analysis

Volumetric imaging of the renal biopsies was performed on a custom-built open-top lightsheet (OTLS) microscope, similar in design to one described in Glaser et al^27^. The microscope has a lateral resolution of 1 µm and axial resolution of 2.7 µm. Images were post-processed using Bigstitcher plugin in FIJI and custom python scripts, yielding large datasets of 500 gigabytes per tissue slice. Approximately 300 glomeruli were manually segmented from 9 different biopsies. Nuclei segmentation was performed in 3D on ∼22% of a biopsy volume using the nuclei model provided in Cellpose 2.0^28^.

### Spatial transcriptomics using digital spatial profiling

A single representative 5 µm slice per condition for each donor was affixed to a SuperFrost Plus charged slide and baked at 60°C for 1 hour. Digital spatial profiling using the Nanostring GeoMx platform was performed as described previously^30^. Briefly, after deparaffinization and heat induced antigen retrieval (Tris EDTA, pH 9), sections were treated for 15 minutes at 37°C with 1µg/ml proteinase K. Slides were then hybridized overnight with the Whole Transcriptome Atlas (WTA) probeset. The next day, slides were washed stringently and labeled with fluorescently labeled antibodies recognizing pan-cytokeratin (532 nm channel), CD45 (594 nm channel), CD31 (647 nm channel) as well as DNA (Syto13, 488 nm channel) (**Fig. 1A**). After loading onto the instrument and scanning, ROI selection was performed, and UV-released barcode probes were collected into a 96-well collection plate. Sequencing libraries were generated using a SeqCode construction kit from Nanostring. The libraries were sequenced using an Illumina NextSeq 6000. Sequenced barcodes were mapped to genes and ROIs using the Nanostring GeoMx NGS Pipeline, version 2.0.21. Two GeoMx Whole Transcriptome Atlas (WTA) experiments were performed on human kidney tissue slices from 8 donors across 2 batches as outlined in **Supplemental Table 3**.

**Figure 1.**
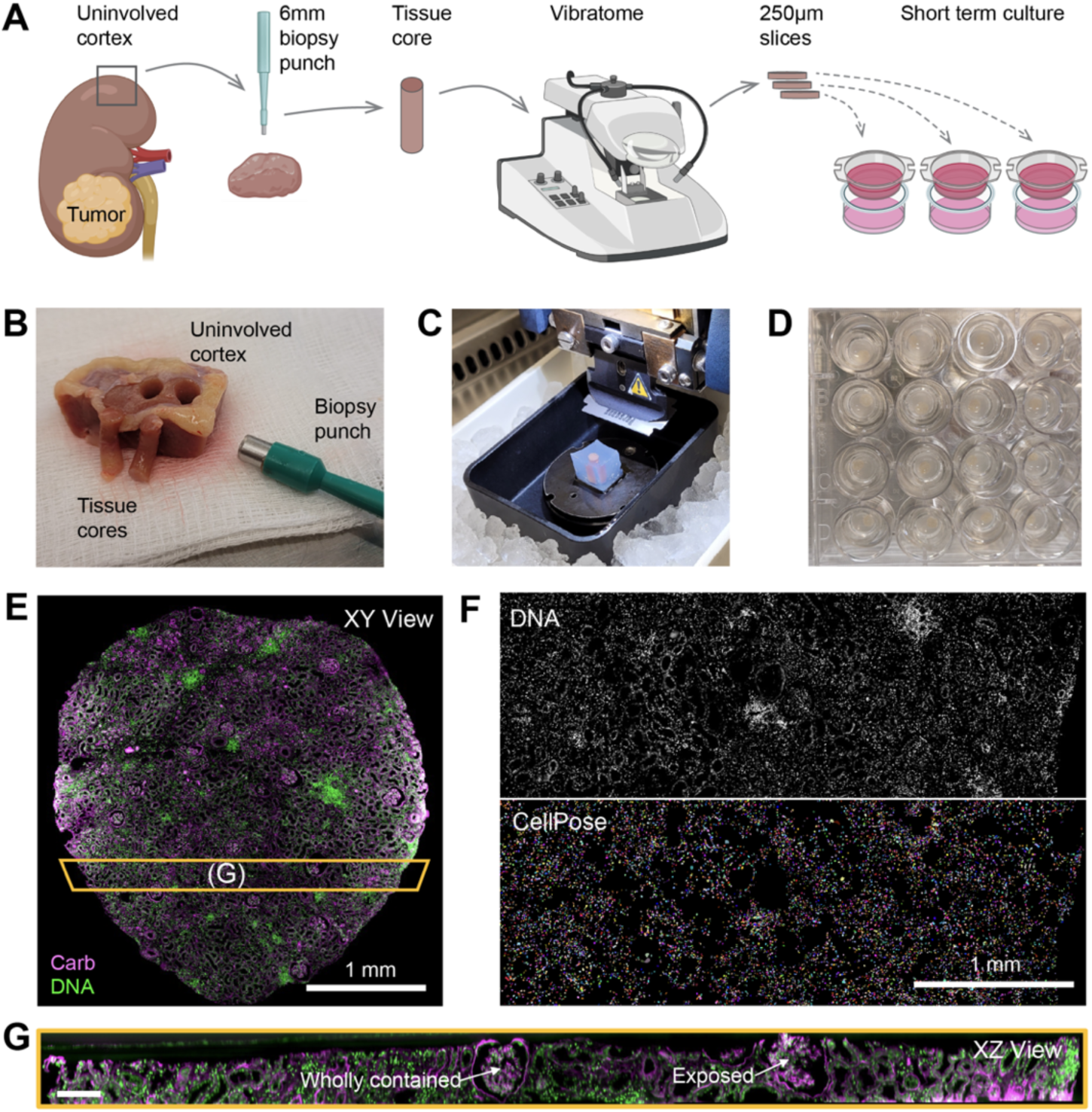
Overview of precision cut kidney slice culture and structural assessment using lightsheet microscopy. A) Schematic representation of precision cut kidney slice culture workflow. B) Photograph of a portion of human kidney cortex and tissue cores generated using a 6 mm biopsy punch. C) Tissue cylinders embedded in low melting temperature agarose and mounted onto the vibratome platform. The collection chamber is packed with ice and shown without collection buffer in order to visualize the agarose block. D) Semi-transparent kidney slices placed in individual transwell inserts in a 24-well plate and incubated with Williams medium without indicator or dyes. E) Lightsheet microscopy overview of kidney tissue slice stained with FLARE protocol. Carbohydrates, purple; DNA, green. F) Portion of tissue slice volume used for 3D nuclear segmentation using Cellpose. Top panel, raw nuclear signal; bottom panel, imputed nuclear contours. G) X-Z projection of portion of slice indicated in panel E.

### Spatial transcriptomics data analysis

Log2-transformed raw GeoMx counts from both batches were combined and quantile-normalized separately for glomerular and tubulointerstitial ROIs with normalization efficacy assessed using PCA^30^. Differential gene expression analysis was performed using *limma* with empirical Bayes moderation including donor as a blocking factor. Genes were considered significant at FDR < 0.05 and |log2FC| > 0.5. To assess culture media effects, a 3-group model (T0, Medium A and Medium B-O2, both at T16) was fit across three donors with pairwise contrasts (A vs T0, B-O2 vs T0, B-O2 vs A) evaluated in both glomerular and tubulointerstitial ROIs. To assess the IFNγ response, per-donor differential expression (IFNγ vs. control, T16, glomerulus) was computed independently across five donors and combined via fixed-effects inverse-variance weighted meta-analysis. Similarly, the effect of the JAK1/2 inhibitor baricitinib^31^ was assessed by two-donor meta-analysis (MTS008, MTS029) for contrasts of Baricitinib vs.

DMSO and IFNγ vs. IFNγ + Baricitinib. Pathway-level analyses used two approaches. First, single-sample GSEA (ssGSEA; *GSVA* package) with MSigDB Hallmark gene sets was used to compute per-ROI enrichment scores and tested for significance across conditions using linear models with Benjamini-Hochberg (B-H) correction. Second, fast preranked GSEA (*fGSEA*) was applied to meta-analysis-derived gene rankings to identify enriched Hallmark pathways outputting normalized enrichment scores with B-H corrected p-values. Temporal expression dynamics over 72 hours (single donor MTS028, B-O2 medium) were characterized by classifying genes into empirically derived trajectory patterns and visualized as z-score-normalized heatmaps. All analyses were performed in *R* v.4.5.1. To assess overlap of gene sets with IFN-regulated pathways, we compared overlap against the union of HALLMARK_INTERFERON_GAMMA_RESPONSE, HALLMARK_INTERFERON_ALPHA_RESPONSE, HALLMARK_IL6_JAK_STAT3_SIGNALING, HALLMARK_IL2_STAT5_SIGNALING, HALLMARK_ALLOGRAFT_REJECTION, HALLMARK_INFLAMMATORY_RESPONSE.

### Data sharing and availability

Spatial transcriptomics data described in this manuscript can be interactively explored at https://faculty.washington.edu/shreeram/GeoMx_Explorer/.

## RESULTS

### Kidney slice protocol and tissue architecture

An overview of the precision cut slice culture approach is shown in **Figure 1A**. Approximately 5 g of macrodissected human kidney cortex was used for most slice culture experiments. A 6 mm biopsy punch was used to produce cylindrical tissue cores (**Figure 1B**). These cores were embedded into low melting temperature agarose for vibratome sectioning (**Figure 1C**). 250 µm thick sections were obtained and placed into transwell inserts in a 24-well plate for culture (**Figure 1D**). Following tissue clearing, fluorescent labeling of nuclei and carbohydrates was performed using the FLARE protocol ^24^ and the entire slice volume was visualized using 3D lightsheet imaging (**Figure 1E**). 3D nuclear segmentation was performed on image stacks from a ∼22% subset of a slice using Cellpose^28^ (**Figure 1F**). This identified 165,579 nuclei which is likely an underestimate since visual inspection showed that approximately 20% of nuclei were not identified due to areas of high autofluorescence and some dense fibrotic clusters where the segmentation failed. Even with this caveat, extrapolating to the entire slice yielded 752,632 nuclei/slice or 107,428 nuclei/mm^3^, which approximated the 143K-250K nuclei/mm^3^ cell/nuclear density in human kidney reported previously^32^. Manual tracing of Bowman’s capsular profiles in z-sections from 8 tissue slices, revealed 32.5±10.8 glomeruli per section (Supplemental Movies 1, 2). This visualization and XZ views showed that on a per slice basis, 81.4±17.8% of glomeruli had a discontinuity in their Bowman’s capsule resulting in direct exposure to the culture medium at the top or the bottom (**Figure 1G**). These experiments showed that precision cut kidney tissue slices yielded structurally intact kidney tissue suitable for subsequent experiments.

### Kidney slice viability and histology in culture

Next, kidney tissue slices were cultured on transwell inserts to assess their viability and morphology over time. In initial experiments, kidney tissue slices were cultured in William’s medium with 5% FBS in 5% CO_2_ in normoxia (Medium A), following published protocols for liver and kidney slice cultures^25^. Transcriptomic characterization of these slices revealed hypoxic and stress signatures, and glycolytic reprogramming (see below). Subsequently, an alternative defined component medium with antioxidants (Medium B) was formulated to prevent oxidative stress and serum-mediated injury, and slices were incubated in carbogen (95% O_2_/5% CO_2_) to limit hypoxia and glycolytic reprogramming. The impact of media and oxygen was assessed in 16-hour tissue slice cultures from 2 donors (MTS023, 024; **Figure 2A**). Tissue viability was measured by quantifying release of the cytosolic enzyme LDH from dead or dying cells into the tissue culture media. While increased O_2_ improved tissue viability of slices cultured in both media, maximum viability was seen when slices were cultured in Medium B with increased oxygen. An initial experiment showed that Medium B-O_2_ preserved approximately 70% viability even up to 72 hours in culture (MTS028; **Figure 2B**). A full 72 hr time course in both media using another donor’s PCKS (MTS045), confirmed the increased viability of tissue slices in Medium B-O_2_ (**Figure 2C**).

**Figure 2.**
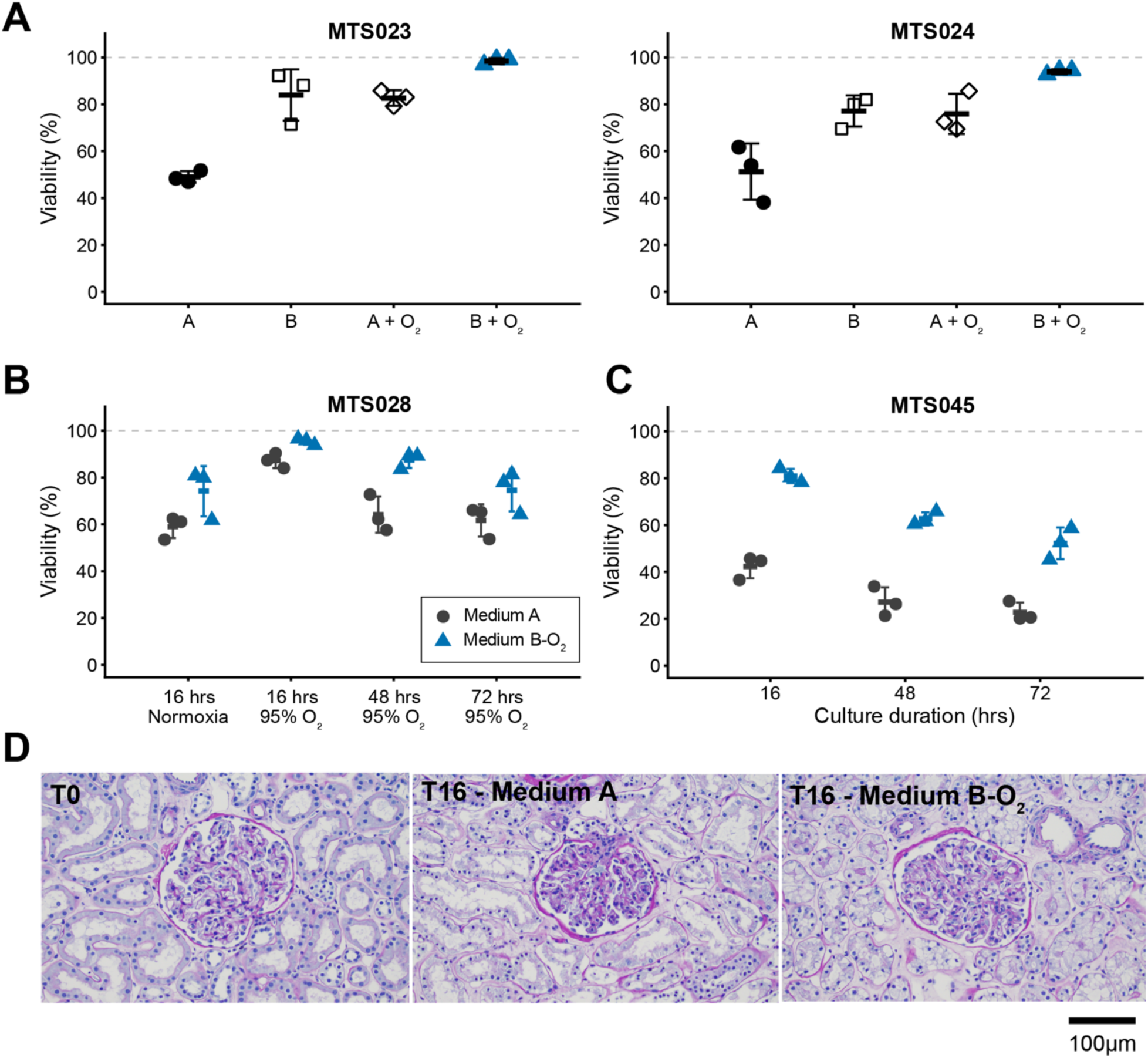
Evaluation of media and oxygenation on kidney slice viability and morphology. A) Viability of triplicate tissue slices after 16 hours of culture (T16) incubated in the indicated medium and oxygenation conditions and assessed using the LDH release assay. Replicate slices lysed using TX-100 were used to determine maximal LDH release. Error bars indicate standard deviations of triplicate slice measurements. B) Initial experiment studying the impact of media and oxygenation on slice viability over 72 hours of culture. C) Repeat experiment studying viability in Medium A and B-O_2_ over 72 hrs of culture. For A-C, bar indicates mean and whiskers represent ±1 standard deviation. Each condition was evaluated with 3 tissue slices per donor. D) PAS-stained sections of donor MTS028’s kidney slices at T0 and after incubation for 16 hrs in medium A or B-O_2_

Examination of PAS-H-stained sections revealed intact nuclear profiles in glomeruli and tubules at 16 hours of culture in Medium A (**Figure 2D**). However, starting at 16 hours (T16), some tubules showed shedding of cells into their lumens. These changes became more widespread with longer durations of culture, with focal necrosis of some tubular segments and loss of nuclear staining with hematoxylin, which was accompanied by an expansion of the interstitium with acellular material. By contrast, tubular epithelial cells showed less detachment and necrosis over 72 hours when cultured in Medium B-O_2_. However, tubular epithelial cells exhibited cytoplasmic vacuolization/hydropic changes that were seen only in the presence of increased O_2_, regardless of media formulation (**Supplemental Figure 1**). These experiments demonstrated that both medium formulation and atmospheric O_2_ conditions had distinct impacts on kidney slice viability and histology.

### Spatial transcriptomics performed on kidney tissue slices

In order to understand tissue responses to culture conditions and stimuli, spatial transcriptomic profiling was performed on tissue slices using the GeoMx digital spatial profiler platform. In this approach, whole transcriptome information is captured from user-defined multicellular regions of interest (ROIs) selected by a trained renal pathologist (S.A.). After quality control, data from only one segment was excluded due to sequencing failure (<1,000 reads) resulting in 464 high-quality segments (343 glomerular, 121 tubulointerstitial). A limit of quantification (LOQ) was calculated using negative control probe signal for each ROI. After filtering for genes detected above the LOQ in ≥10% of segments within at least one biological group, 9,979 of the 18,676 genes in the WTA (53.4%) were retained for downstream analysis. Glomerular and tubulointerstitial ROIs were quantile normalized separately (**Supplemental Figure 2**). Even without formal batch correction, principal component analysis (PCA) demonstrated a reduction of batch effect (PC1 variance reduced from 76-85% to ∼10%) (**Supplemental Figure 3**). In subsequent analyses, two strategies were employed to further reduce the impact of batch effect. First, differential expression calculations were performed as within-donor contrasts. Second, cross-donor comparisons utilized inverse-variance weighted fixed-effects meta-analysis, which combines within-donor effect sizes (i.e., fold-changes) without requiring absolute expression levels to be comparable across batches.

### Global transcriptomic responses of kidney tissue slices to different culture conditions

Spatial transcriptomics data were generated from three donors (MTS023, MTS024, MTS028) with slices cultured for 16 hours (T16) in either Medium A or B-O_2_. Compared to freshly cut tissue (T0), culturing slices for 16 hrs in both media conditions induced expression changes in ∼700 genes in both glomerular and tubulointerstitial ROIs. Compared to T0, glomeruli from slices cultured in Medium A demonstrated a strong hypoxia signal and increased expression of multiple HIF target genes (e.g., *BNIP3, BNIP3L, NDRG, HILPDA, VEGFA, EGLN3, PDK1, SLC2A1 and SLC2A3*), induction of glycolytic pathway genes (e.g., *HK2, PFKFB3, PGK1, GLUT1, GLUT3, LDHA and ENO1*) and integrated stress responses with induction of genes in endoplasmic reticulum (ER), translational, nutrient and hypoxia stress pathways (e.g., *DDIT4, DDIT3, PPP1R15A, HSPA1A* and *HSPA1B*). Glomeruli also downregulated podocyte-specific genes (e.g., *SYNPO, NPHS2, PTPRQ, PLA2R1* and *TRPC6*) and endothelial markers (e.g., *EMCN, EHD3, CD34, PTPRB, SOX17*, and *ERG*). Compared to T0, glomeruli cultured in Medium B-O_2_ also showed activation of hypoxia, inflammatory and glycolytic pathway genes, but these increases were blunted compared to Medium A. Notably, slices incubated in Medium B-O_2_ showed induction of genes involved in endothelial cell maintenance, growth, and remodeling (e.g., *PECAM1, KDR, CLDN5, RAMP2, EMCN, SHB, KDR, NRP1* and *ENG*).

In summary, culturing slices in Medium A for 16 hrs induced a robust transcriptional response characterized by activation of hypoxia, inflammatory, and glycolytic pathways accompanied by suppression of endothelial differentiation markers. These data suggested that short-term culture in Medium A triggers hypoxic stress, endothelial dysfunction, metabolic adaptation, and early tissue remodeling in human glomeruli. By comparison, culturing slices in Medium B-O_2_ for 16 hrs showed continued activation of inflammatory pathways, but with reduced hypoxic stress and glycolytic reprogramming, and with preservation of endothelial cell differentiation. Together, these data indicated that short-term culture in Medium B-O_2_ blunted hypoxic stress, endothelial dysfunction, and metabolic adaptation relative to Medium A. Analysis of the DEGs in the tubulointerstitium between T0 and Medium A or Medium B-O_2_ further supported these conclusions. The tubulointerstitial ROIs from slices cultured in Medium A showed severe epithelial stress and a shift towards anaerobic metabolism as evidenced by strong heat shock, ER stress, and hypoxia responses with glycolytic rewiring and tubular epithelial cell dedifferentiation. Culturing the slices in Medium B-O_2_ partially reversed these stress responses in tubulointerstitial ROIs, but increased expression of genes involved in inflammatory and remodeling pathways (**Supplementary Table 4**).

Direct comparison of Medium A and Medium B-O2, reinforced the above results and identified additional differences. Each medium elicited unique expression changes with145 DEGs specific to glomeruli and 257 DEGs to tubulointerstitium (**Figure 3E, F**). Single-sample gene set enrichment analysis (ssGSEA) using Hallmark pathways revealed many shared pathways that were induced by both media conditions in glomeruli, including TNFA_SIGNALING_VIA_NFKB, MYC_TARGETS and INFLAMMATORY_RESPONSE (**Figure 4A, B**, **Supplemental Table 5**).

**Figure 3.**
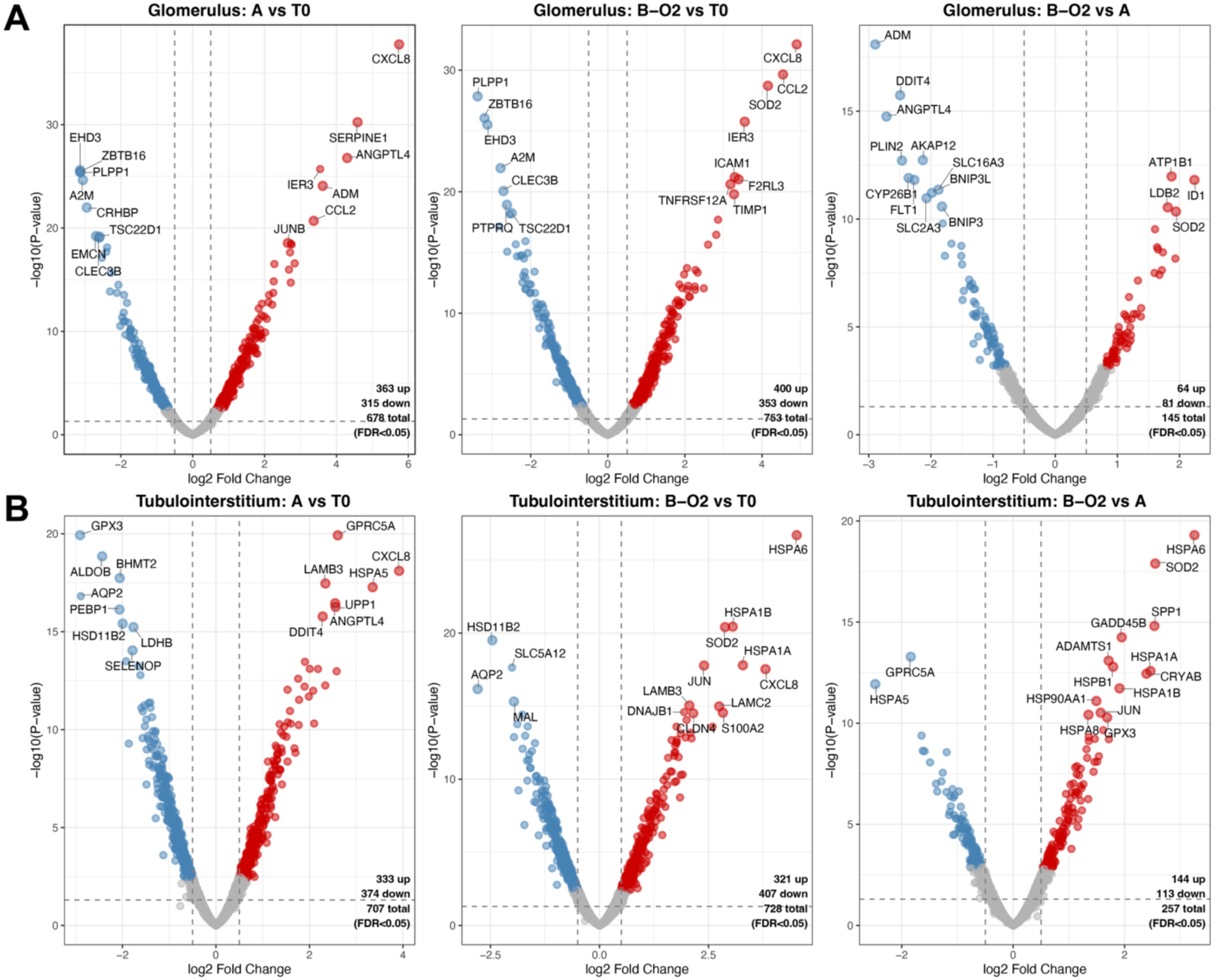
Spatial transcriptomic characterization of glomerular and tubulointerstitial response to 16-hour culture. Volcano plot of differentially expressed genes in A) glomerular and B) tubulointerstitial ROIs after 16 hours of culture in medium A or B-O_2_.

**Figure 4.**
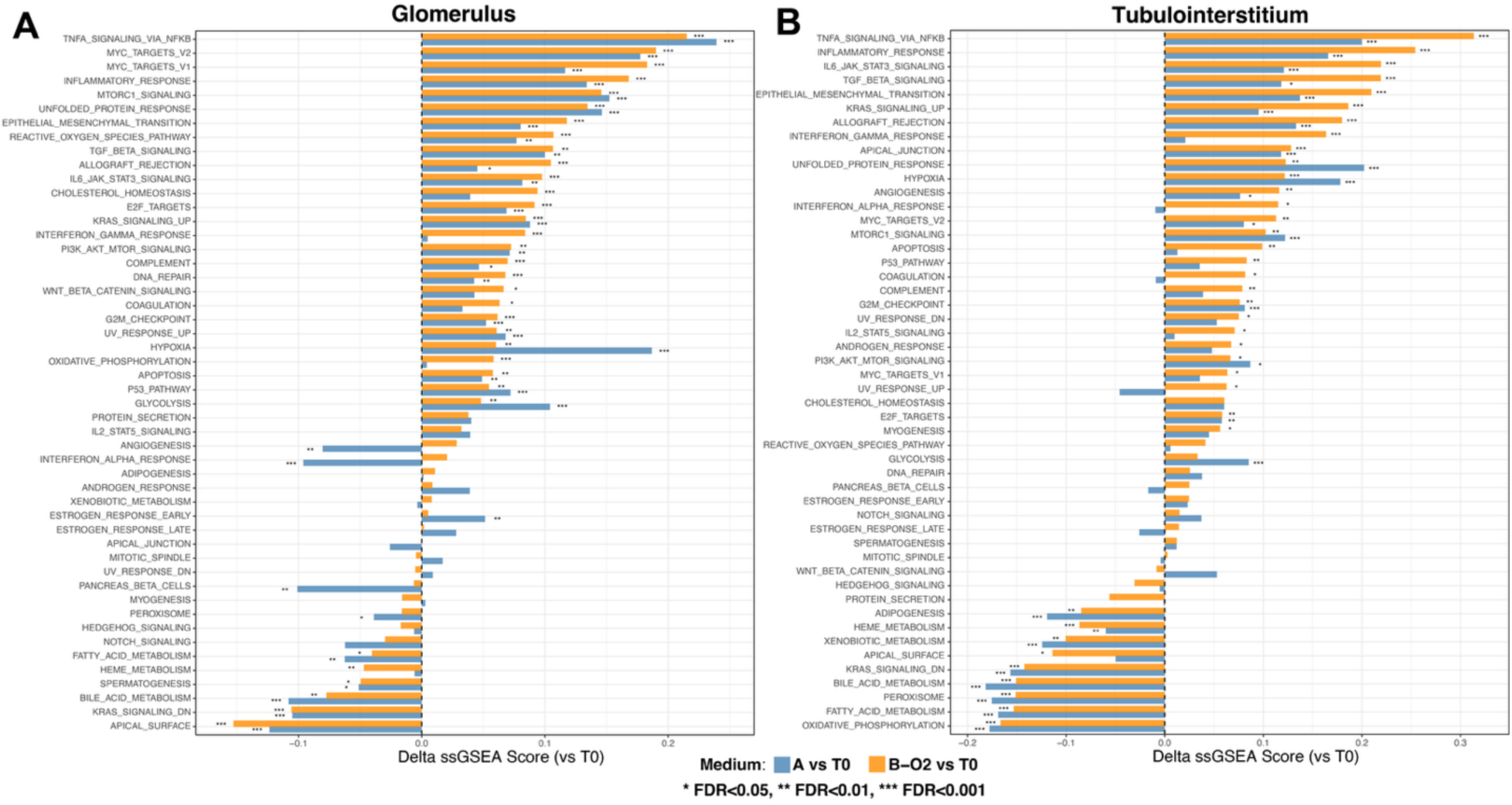
Hallmark pathway changes in glomeruli and tubulointerstitium after 16-hour culture in different media conditions. The difference in ssGSEA (delta ssGSEA score) for Hallmark pathways in A) glomerular and B) tubulointerstitial ROIs when cultured in medium A or B-O2 (compared to T0 tissue slices). * FDR< 0.05, ** FDR<0.01, *** FDR<0.001.

While both media conditions induced HYPOXIA and GLYCOLYSIS pathway signatures, this was more pronounced in Medium A. For example, hypoxia-inducible genes *ANGPTL4*, *ADM* and *DDIT4*, were elevated in glomeruli cultured in Medium A compared to those cultured in Medium B-O_2_. Inflammatory pathways were induced in both media, with Medium B-O_2_ showing more pronounced induction of interferon-associated signatures (e.g., ALLOGRAFT_REJECTION) and in particular, the INTERFERON_GAMMA_RESPONSE signature, which was not induced in Medium A. The relative increase in inflammatory signatures in Medium B-O_2_ coincided with a stronger REACTIVE_OXYGEN_SPECIES_PATHWAY signature, suggesting that reactive oxygen species (ROS) generated in the high O_2_ atmosphere may have driven the inflammatory phenotype in Medium B-O_2_ cultured slices. Glomeruli cultured in Medium A also had lower ssGSEA scores for ANGIOGENESIS, reflecting the selective loss of expression of endothelial maintenance and growth in Medium A.

The tubulointerstitium had some overlap with glomeruli but also had distinct pathway differences in both media. Inflammatory signatures (INTERFERON _GAMMA_RESPONSE, INTEFERON_ALPHA_RESPONSE) were higher in the tubulointerstitium compared to glomeruli, perhaps reflecting the increased inflammatory cells (e.g., tissue-resident macrophages) in this compartment, and similar to glomeruli these signatures were increased in Medium B-O_2_ compared to A. In glomeruli. Medium B-O_2_ showed better preservation of the OXIDATIVE_PHOSPHORYLATION, FATTY_ACID_METABOLISM, and PEROXISOME signatures; whereas culture in both media suppressed these signatures in tubulointerstitium.

These results showed that increasing ambient O_2_ during culture effectively suppressed the hypoxia and glycolysis-associated signatures. However, culturing slices in Medium B-O_2_ increased interferon, inflammatory, and reactive oxygen species associated signatures. Importantly, compartment-specific analysis of the slice culture conditions revealed differences between the glomeruli and tubulointerstitial compartments, such as the greater angiogenesis and oxidative phosphorylation signatures in Medium B-O_2_ culture condition in glomeruli compared to the tubulointerstitium. In addition, many pathways remained relatively unperturbed by slice culture. The analysis shows the strength of employing spatial transcriptomic analysis to understand the biological strengths and weaknesses of the experimental system for specific tissue compartments, and how to use this knowledge to select conditions or design new configurations that are optimal for the experimental question.

### Global transcriptomic responses of kidney tissue slices over 72 hours of culture

Next, spatially localized gene expression changes were studied for glomerular and tubulointerstitial ROIs from one donor (MTS028) over 72 hours of culture in Medium B-O_2_, since that culture condition maximized slice viability. Three complementary approaches were combined to identify dynamically expressed genes. First, ANOVA was used to identify genes that were variably expressed at any of the four timepoints (T0, T16, T48, T72). Second, a linear regression was used to identify genes with a non-zero slope across the time course. Third, the expression of genes at each timepoint was compared to the other timepoints to reveal genes that had a sharp peak or dip at any given timepoint. The union of genes with FDR <0.05 by any of these strategies was then ranked by the ANOVA F-statistic, with the top 100 genes for glomerulus and tubulointerstitium plotted in **Figure 5** and listed in **Supplemental Table 6**. This approach identified distinct temporal expression patterns as well as differences between glomeruli and tubulointerstitium. In glomeruli, some genes important for endothelial cell (*EHD3*), mesangial cell (*PTPRQ*), and podocyte (*CDKN1C*) function or identity were rapidly downregulated by 16 hrs of culture. Other genes showed a more gradual decline in expression over the 72 hr time course. Some pro-inflammatory genes were rapidly induced at 16 hrs of culture (*CCL2*, *CXCL8*). Other pro-inflammatory genes, growth factors, and several extracellular matrix-related genes showed sustained increases in expression over the 72 hrs of culture (*IL32*, *PGF*, *COL4A1*/2, *FN1*, *LAMA4*, *ANXA2*). In the tubulointerstitium, some solute transporters showed gradual decline in expression over time (*SLC16A9*, *SLC5A12*, *SLC3A1*, *SLC22A8*). The AP-1 response pathway components (*FOS*, *JUN*, *JUNB*), and cellular stress response genes (*HSPA1A/B*, *DNAJB1*, *GADD45B*) were strongly induced at the 16-hour timepoint. Similar to glomeruli, within the tubulointerstitium proinflammatory and extracellular matrix-related genes increased in expression over time in culture (e.g., *IL6*, *CXCL8*, *KRT7*, *BGN*, *VIM*). Since protein levels may not match gene expression levels, the slice culture system may still be suitable for the study of biological processes where protein abundance remains relatively unchanged over the incubation period, as was the case for NPHS1 and SLC6A19 (**Supplemental Figure 4**).

**Figure 5.**
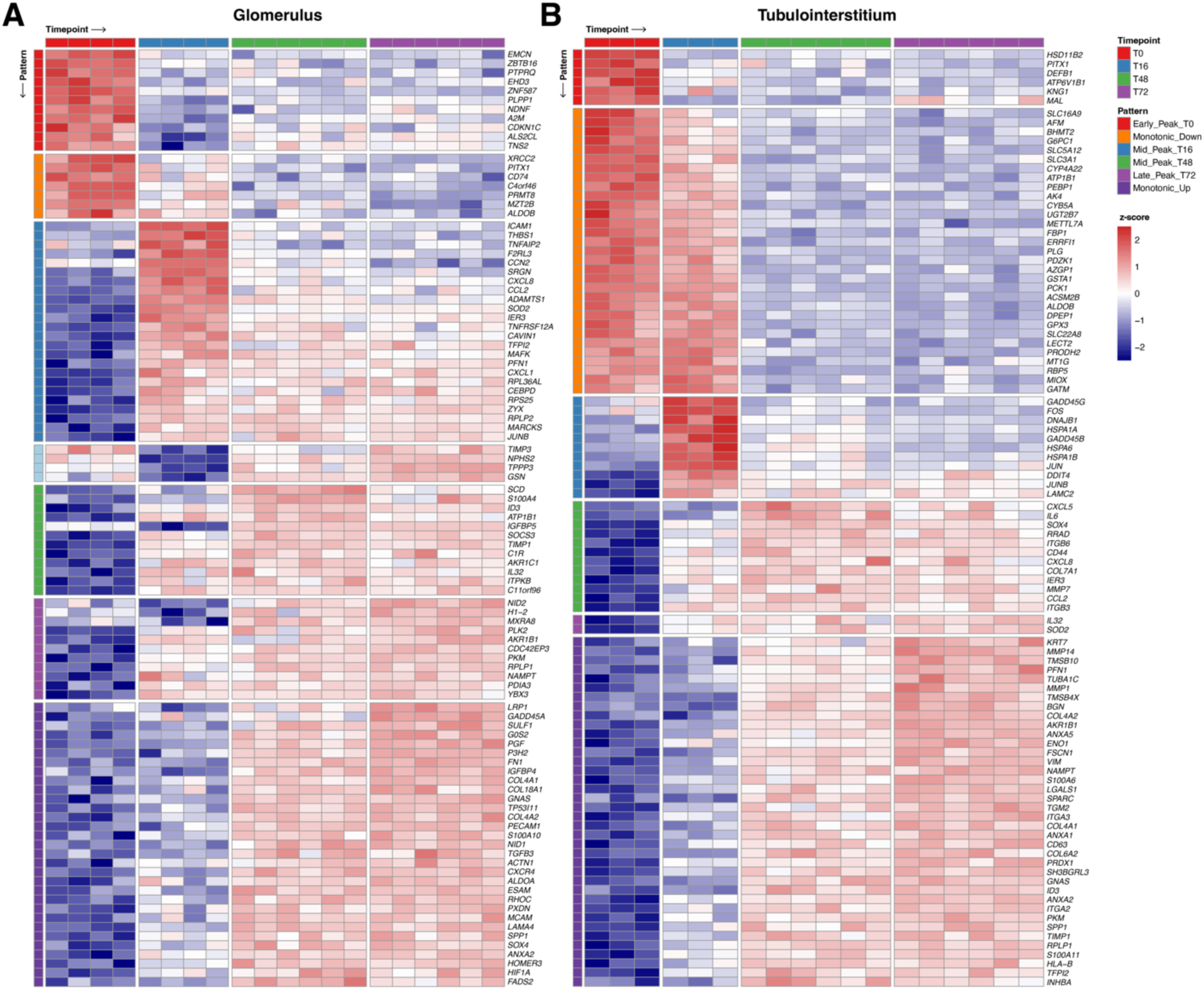
Temporal expression patterns of 100 top-ranked differentially expressed genes. Relative expression (z-score normalized) of 100 top-ranked differentially expressed genes in A) glomerular and B) tubulointerstitial ROIs displaying different patterns of expression changes over 72 hours in culture.

These experiments showed that the glomeruli and tubulointerstitium exhibited distinct temporal patterns of gene expression in *in vitro* slice culture over 72 hours.

### APOL1 as a marker of biological responsiveness of precision cut kidney tissue slices

Next, we explored the biological responsiveness of *APOL1*, a known IFNγ-regulated gene important for kidney disease, in the PCKS system. PCKS were treated with IFNγ for 16 hrs and baricitinib, a JAK1/2 inhibitor that is approved to suppress IFNγ signaling in rheumatoid arthritis, was applied to slices to test its ability to block IFNγ-induced *APOL1* expression, as assessed by spatial transcriptomics. *APOL1* was consistently induced in glomeruli, though with some donor-to-donor variability (**Figure 6A**). The lower relative induction in MTS029 was due to higher baseline IFN pathways in the vehicle-only treated slices when cultured in Medium B-O_2_ (**Figure 6B**). Baricitinib effectively suppressed IFNγ-induced *APOL1* expression, even below the vehicle-only levels, irrespective of incubation media (**Supplemental Figure 5A**). These spatial transcriptomic changes in *APOL1* expression were orthogonally validated at the whole slice level using QPCR (**Supplemental Figure 5B**), RNA-ISH and immunohistochemistry (**Figure 6C**). These studies showed that precision cut kidney tissue slices could be used to study the regulation and therapeutic modulation of a kidney-relevant gene within its native anatomic context.

**Figure 6.**
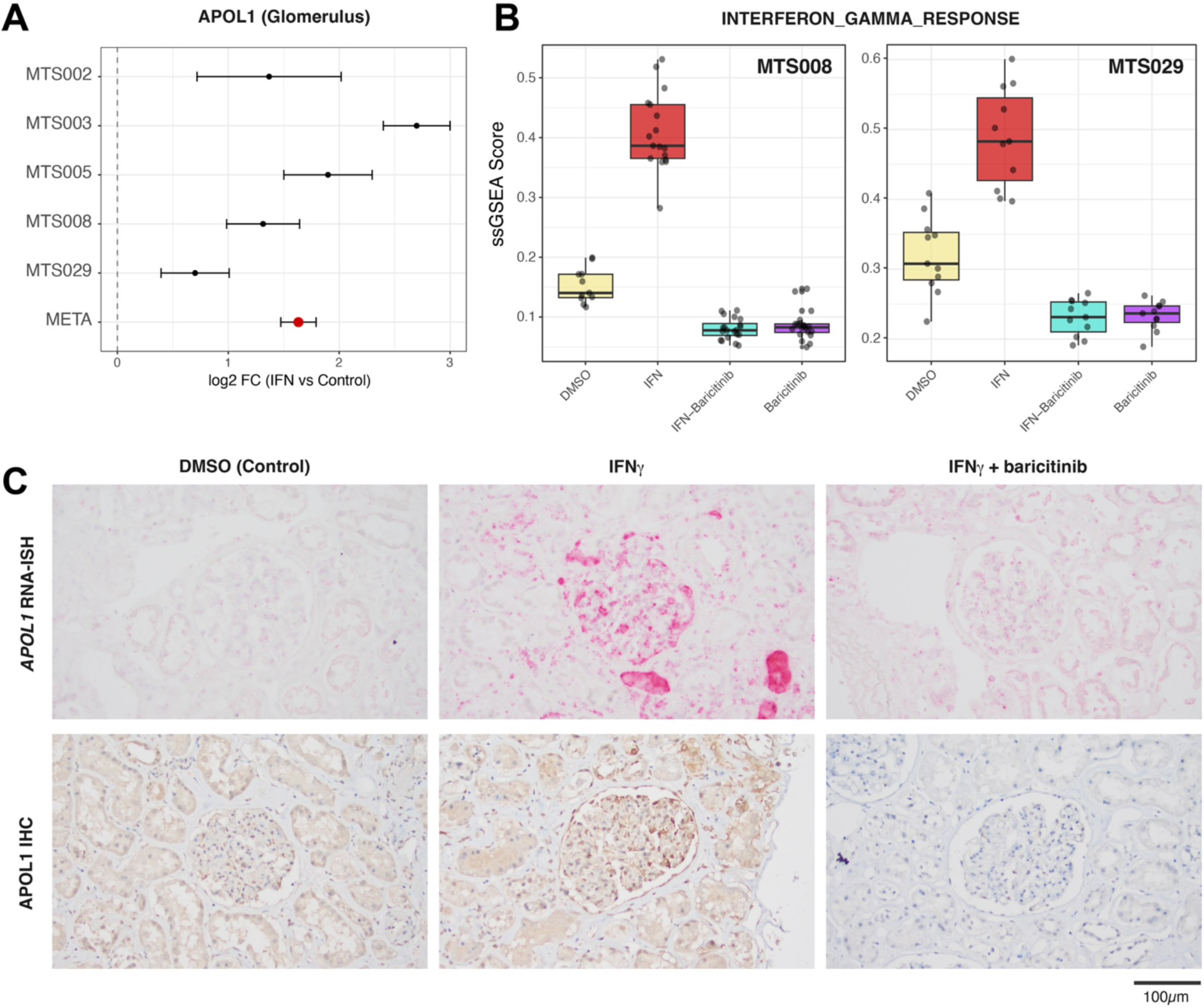
Glomerular APOL1 responses to IFNγ stimulation and baricitinib. A) Forest plot of glomerular APOL1 expression induced by IFNγ across donors. B) ssGSEA of the Hallmark interferon gamma response pathway in glomeruli from donors MTS008 and MTS029. C) APOL1 mRNA (RNA-ISH, top row) and protein expression (IHC, bottom row) in glomeruli from donor MTS008 PCKS with the indicated treatments.

### Dissection of drug mechanisms in precision cut kidney tissue slices

Next, the utility of PCKS to dissect mechanisms of drug action was explored in depth. Initial experiments had utilized slices cultured in Medium A (donors MTS002, 003, 005, 008) and a later experiment had utilized slices cultured in Medium B-O_2_ (MTS029). Fixed-effects inverse-variance weighted meta-analysis across all five donors identified 2,124 genes (903 upregulated, 1,221 downregulated) with significant expression changes in IFNγ-treated glomeruli vs.

PBS/vehicle only controls (FDR < 0.05). 903 of these differentially expressed genes (426 upregulated, 477 downregulated) showed the same directionality of expression change across all 5 donors (**Supplemental Table 7**). Gene set enrichment analysis (**Figure 7A**) on the ranked gene list from glomeruli confirmed HALLMARK_INTERFERON _GAMMA_RESPONSE (Normalized Expression Score, NES = 3.54, padj < 10⁻⁴⁹) and HALLMARK_INTERFERON_ALPHA_RESPONSE (NES = 3.33, padj < 10⁻⁴⁴) as the most strongly enriched pathways. Suppressed pathways included MYC targets V1/V2, oxidative phosphorylation, and TGF-beta signaling. Comparing IFNγ treated to IFN+baricitinib glomerular ROIs, the role of interferon signaling pathways was magnified due to suppression of baseline interferon pathways (**Supplemental Figure 6**). Gene expression data from tubulointerstitial ROIs were also generated from donor MTS029. However, the lower number of ROIs in this comparison (4 IFN vs. 4 DMSO) limited the power to detect expression differences induced by IFNγ, though interferon-related Hallmark pathways still showed the largest changes (**Supplemental Figure 7**).

**Figure 7.**
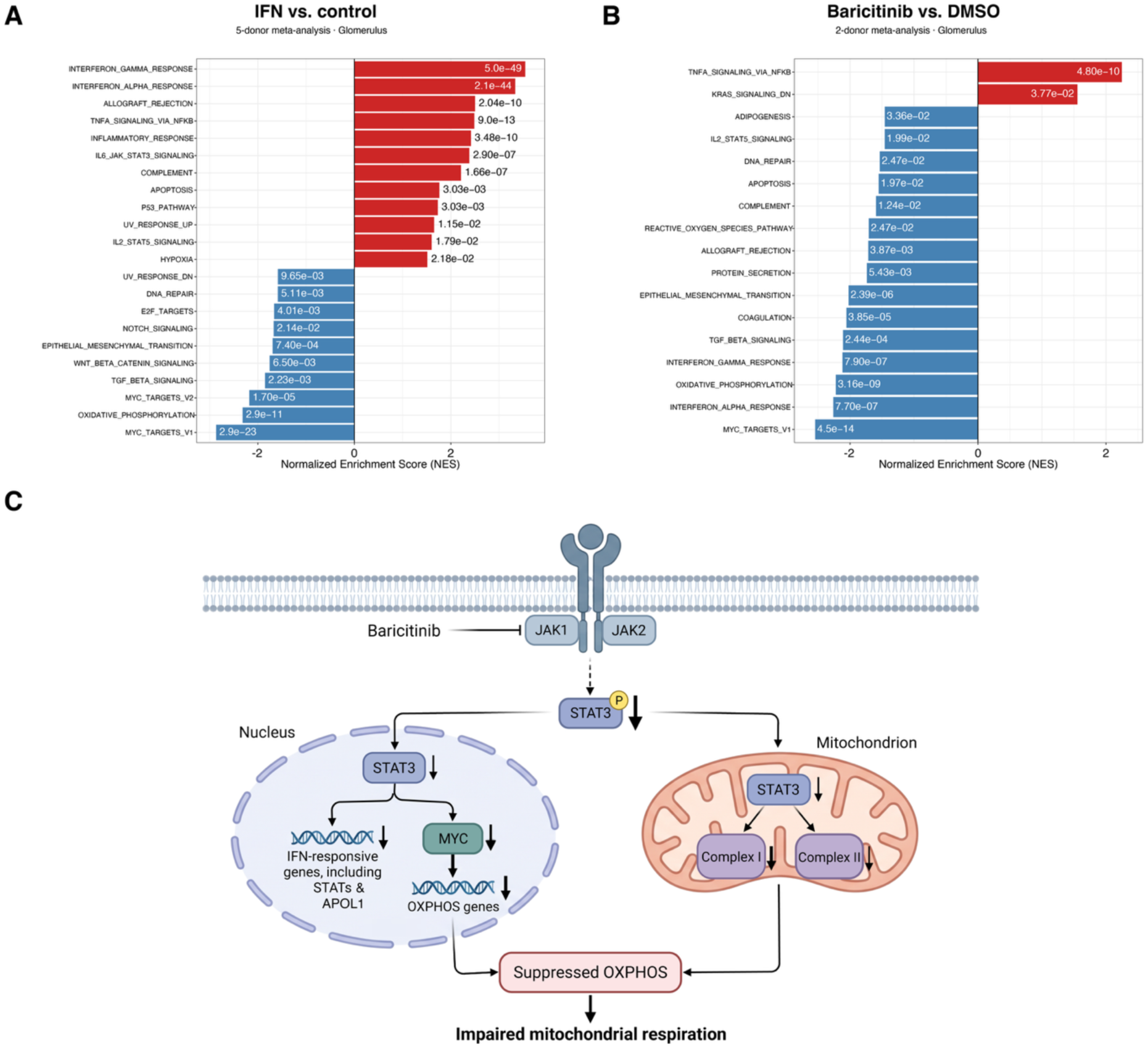
Glomerular pathway responses to IFNg and baricitinib. A) fGSEA of significantly enriched pathways following 16 hours of IFNg exposure compared to control. B) fGSEA of significantly enriched pathways following 16 hours of baricitinib exposure compared to vehicle-only control. C) Proposed mechanism linking baricitinib-mediated JAK1/2 inhibition to suppression of IFN-responsive and STAT3-MYC-associated oxidative phosphorylation pathways. Created in BioRender. Akilesh, S. (2026) https://BioRender.com/3gy9k7r.

Comparing bariticitinib-treated glomerular ROIs to vehicle-only (DMSO) control ROIs from donors MTS008 (Medium A) and MTS029 (Medium B-O2) allowed for dissection of pathways modulated by baricitinib in human kidney tissues. Meta-analysis revealed 1,814 significant DEGs (863 upregulated, 951 downregulated) with 1,712 of these (808 upregulated, 904 downregulated) showing the same directionality of expression change across both donors (94.4% concordance). The full list of baricitinib-associated DEGs is provided in **Supplemental Table 8**. Gene set enrichment analysis (**Figure 7B**) confirmed that baricitinib suppressed HALLMARK_INTERFERON_GAMMA_RESPONSE (NES = −2.12, padj < 10⁻⁷) and HALLMARK_INTERFERON_ALPHA_RESPONSE (NES = −2.26, padj < 10⁻⁷), consistent with potent inhibition of basal JAK1/2-mediated interferon signaling even in the absence of exogenous IFNγ. However, other suppressed pathways included MYC_TARGETS_V1 (NES = −2.55, padj = 4.5 × 10⁻¹⁴), OXIDATIVE_PHOSPHORYLATION (NES = −2.22, padj = 3.2 × 10⁻⁹), TGF_BETA_SIGNALING (NES = −2.11, padj = 2.4 × 10⁻⁴), EPITHELIAL_MESENCHYMAL_TRANSITION (NES = −2.02, padj = 2.4 × 10⁻⁶), and COAGULATION (NES = −2.06, padj = 3.9 × 10⁻⁵), while TNFA_SIGNALING_VIA_NFKB (NES = 2.25, padj = 4.8 × 10⁻¹⁰) and KRAS_SIGNALING_DN (NES = 1.55, padj = 3.8 × 10⁻²) were the only significantly induced pathways (**Figure 7B, Supplemental Table 9**). Hallmark pathways can have redundant representations of genes, but the 2 most significantly downregulated pathways, MYC_TARGETS_V1 (62 genes, 59 unique, 95.2%; *MYC*, *PSMB2* and *NME1* overlapped) and OXIDATIVE_PHOSPHORYLATION (44 genes, 44 unique, 100%) had minimal overlap with IFN-regulated Hallmark pathway gene sets. 100% (44/44) of OXIDATIVE_PHOSPHORYLATION pathway genes and 95.2% (59/62; *PSMC4*, *RPS2*, *SRSF2* were discordant) of MYC_TARGETS_V1 pathway genes showed the same directionality of expression change in both donors despite differences in culture conditions and baricitinib concentrations. The majority of concordantly regulated genes assigned to both of these pathways had decreased expression with baricitinib treatment, 53/59 for MYC_TARGETS_V1 and 41/44 for OXIDATIVE_PHOSPHORYLATION. Glomeruli and tubulointerstitial compartments showed overlapping, but distinct responses to IFNγ stimulation (**Supplemental Table 10**). The ability to use control and baricitinib-only slices from the same donor also enabled dissection of signaling pathways affected by baricitinib independent of IFNγ stimulation. These results demonstrate that PCKS can capture both expected and unanticipated transcriptional effects of targeted therapies within the native tissue microenvironment.

## DISCUSSION

Precision cut kidney slice cultures enable the study of mature human kidney cell types in their native anatomic context. Mature cellular gene expression signatures are stronger in nephrectomy tissues compared to organoid model systems^33^. Standard imaging approaches can also be applied to kidney slices, as well as 3D imaging modalities, such as light sheet microscopy. Stabilization of experimental kidney slices in standard formats (fresh frozen, formalin-fixed paraffin embedded), permits sophisticated, hi-plex analysis with emerging spatial -omics platforms, as we show here using spatial transcriptomics. These unbiased molecular approaches can calibrate tissue responses to culture conditions such as media composition and oxygenation. Understanding the impact of these will influence the choice of culture conditions to optimize for the biological question being studied. These approaches will also enable further data-driven optimization of precision cut kidney slice cultures. While we do not employ them here, dissociative approaches (single cell -omics) could also be applied to slice cultures. These approaches can also readily be applied to studying glomerular disease mechanisms in slice cultures of genetically modified model organisms. This would represent an animal-sparing approach, since an entire experiment with replicates and controls could be executed from tissue slices obtained from a single animal.

As we show here, the combination of kidney slice cultures and spatial transcriptomics can be used to decipher the kidney intrinsic responses to IFNγ and the effect of the JAK1/2 inhibitor baricitinib in the kidney. Using kidney slice cultures and spatial transcriptomics, we have defined the global changes in gene expression that are induced by IFNγ in glomeruli. Using slice cultures, we demonstrate that baricitinib, a potent JAK1/JAK2 kinase inhibitor^31^, is able to effectively suppresses IFN-regulated pathways. However, baricitinib treatment also altered other pathways not clearly regulated by exogenous IFNγ. The suppression of MYC and Oxidative Phosphorylation (OXPHOS) pathways by baricitinib are not typically associated with JAK1/2 inhibition; however, these effects are likely on-target but potentially undesirable consequences of JAK-STAT pathway inhibition. JAK1 and JAK2 signal via STAT1/2/3 to activate inflammatory cytokine and interferon signaling pathways^34–37^. The STATs themselves are interferon-stimulated genes and can reinforce their own expression through positive autoregulatory loops^38, 39^.

Consistent with this, both *STAT1* and *STAT3* were downregulated with baricitinib treatment. STAT3 is a known positive regulator of MYC transcription^40, 41^, potentially explaining the reduced expression of *MYC* and 53 of its downstream targets in baricitinib-treated glomerular ROIs.

MYC in turn directly regulates expression of nuclear-encoded mitochondrial genes^42, 43^, providing a mechanistic link to the broad suppression of OXPHOS genes. This transcriptional effect may be compounded by loss of non-canonical mitochondrial STAT3, which directly supports Complex I and Complex II enzymatic activity independent of transcription^44^. Taken together, these findings suggest a signaling cascade: baricitinib ⊣ JAK1/2 → ↓STAT3 → ↓MYC → ↓nuclear-encoded OXPHOS genes, that connects JAK1/2 inhibition to suppression of mitochondrial function (**Figure 7C**). These observations may explain some of the known side effects of baricitinib in non-kidney tissues. Impairment of MYC and its downstream target genes, including key nucleotide biosynthesis genes *IMPDH2*, *NME1*, and *CTPS1*, may explain severe leukopenia that is seen in some patients. IMPDH1/2 are targets of the purine synthesis inhibitor, mycophenolate mofetil, which can also cause leukopenia. Inhibition of mitochondrial function via MYC and STAT3, may underlie reports of muscle damage and creatine phosphokinase (CPK) release^45, 46^. While it is the therapeutic goal, suppression of interferon-regulated pathways would also be expected to increase susceptibility to infections, as is also seen in patients treated with baricitinib. With respect to its impact on the kidney, baricitinib has been associated with dose-dependent and temporary increases in serum creatinine, a marker of kidney injury^46^. However, these effects appear to be mild and self-limiting and have not led to stoppage of drug use. Still, our experiments suggest that baricitinib can suppress MYC and OXPHOS pathways in glomeruli. It is unclear how this alteration of glomerular expression profiles would impact kidney function or toxicity and therefore, further study of baricitinib-mediated glomerular injury mechanisms may be warranted. More broadly, these results demonstrate that precision cut kidney tissue slices combined with spatial transcriptomics can reveal tissue compartment-specific drug mechanisms.

Even with these advantages, it is important to consider the shortcomings of kidney slice cultures. Not all investigators may have access to human nephrectomy tissues, which requires an institutionally approved process to obtain informed consent, minimize risk to patients, protect privacy and confidentiality, operate under independent ethical oversight, and follow rigorous quality standards to use specimens only in ways that respect donor rights, legal requirements, and the public trust. Patients undergoing nephrectomy for the removal of cancer will tend to skew older, male, and are likely to have comorbidities such as smoking history, diabetes and hypertension, which could impact the questions being studied. Compared to experimental animal models, donor-to-donor variability is inherent to human slice cultures, though understanding this variation is likely to be important in treating patients with kidney disease.

Even with this variability, consistent tissue responses may still be apparent, as we observed for viability assays and transcriptional responses to IFNγ stimulation and baricitinib treatment. Perhaps the most important consideration is that the tissue will be undergoing continuous injury as soon as it is extracted from the body. Transcriptomic analysis of human kidney slice cultures has shown that pro-fibrotic pathways are activated during *in vitro* culture^47^. Even with rapid processing of nephrectomy tissues, injury signatures will be present when comparing them to living donor biopsy tissues^48^. Modified culture conditions will likely be required to address experimental questions that require long term incubation. Lastly, genetic manipulation and introduction of fluorescent reporters, such as been shown for organoid cultures^49^, will be more difficult in slice cultures due to their size and heterogeneity, and limited replicative capacity.

In summary, we demonstrate the feasibility of the slice culture paradigm for multiple donors using cytokine treatment and small molecule/drug-mediated inhibition. Genes representing fundamental cellular processes are preserved in short-term culture, though there are differences between glomerular and tubulointerstitial compartments. Short term precision cut kidney slice culture therefore represents a biologically responsive and physiologically relevant tissue platform to study mechanisms of glomerular disease.

## ACKNOWLEDGMENTS

We would like to thank the NWBioSpecimen project coordinators and pathology assistants, especially Piper Driskell and Aldous Subang, for consenting patients and procuring deidentified fresh kidney tissues for these experiments. We would like to thank Dr. Raymond Yeung (UW Department of Surgery) and his laboratory members for generously training our group on use of a vibratome to generate precision cut kidney slices. The UW Spatial Biology Core Facility (a.k.a., <u>M</u>olecular <u>An</u>notation and <u>T</u>issue Interrogation <u>S</u>ervice, MANTIS) was supported in part by the UW Department of Laboratory Medicine and Pathology. S.A. and K.D.S. were supported in part by R01 DK130386. Experiments were also supported by the True-Brown Endowed Chair in Laboratory Medicine and Pathology to S.A. C.P. was supported by a Washington Research Foundation Postdoctoral Fellowship. P.V. was supported by a NIDDK NCOR-KUH TL1DK143270 traineeship. J.C.V. was supported by NIH grants DK148406 and GM163162.

---

**Supplemental Figure 1.**
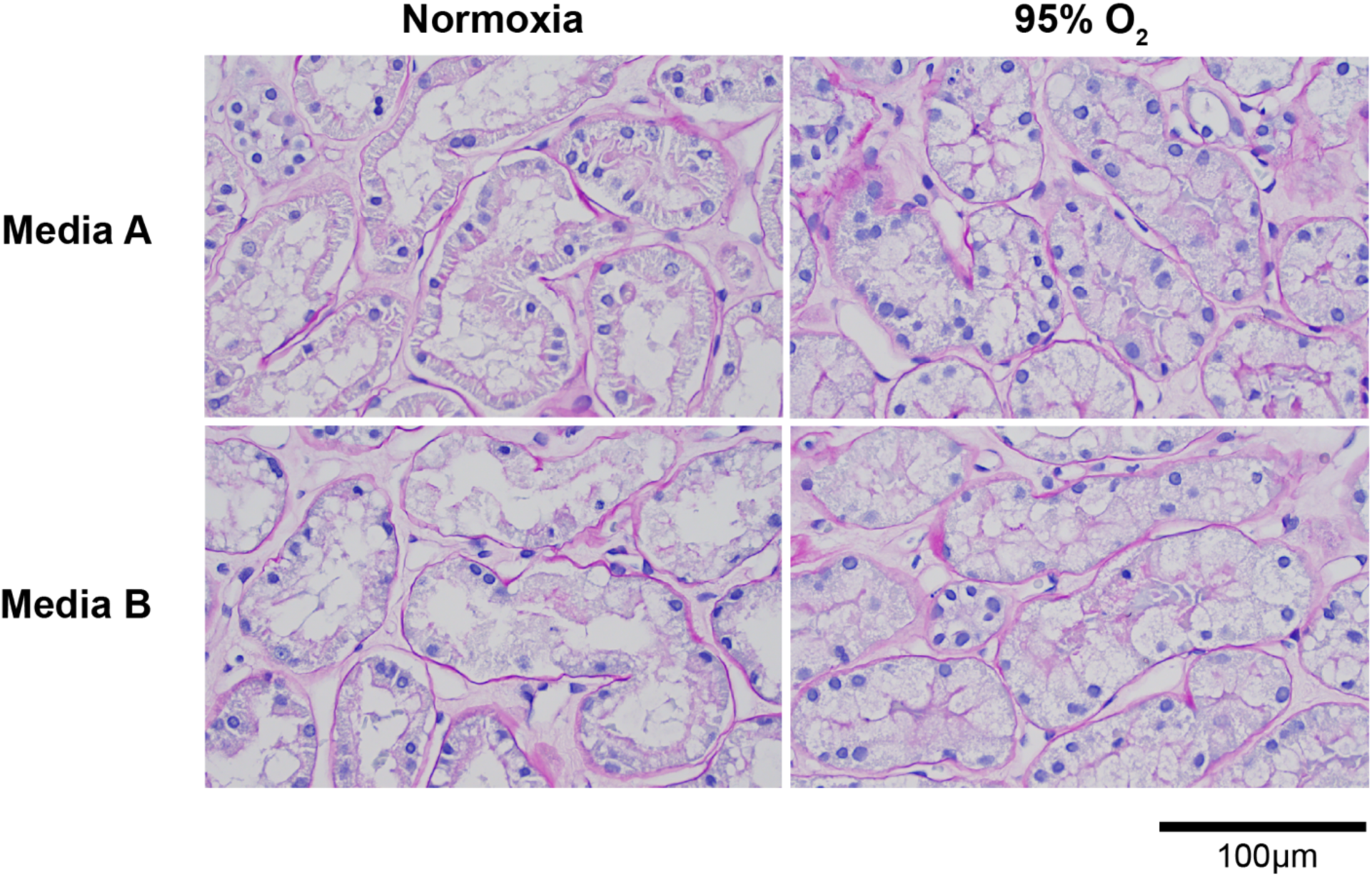
Impact of media and oxygen on tubular histology at 16hrs of culture.

**Supplemental Figure 2.**
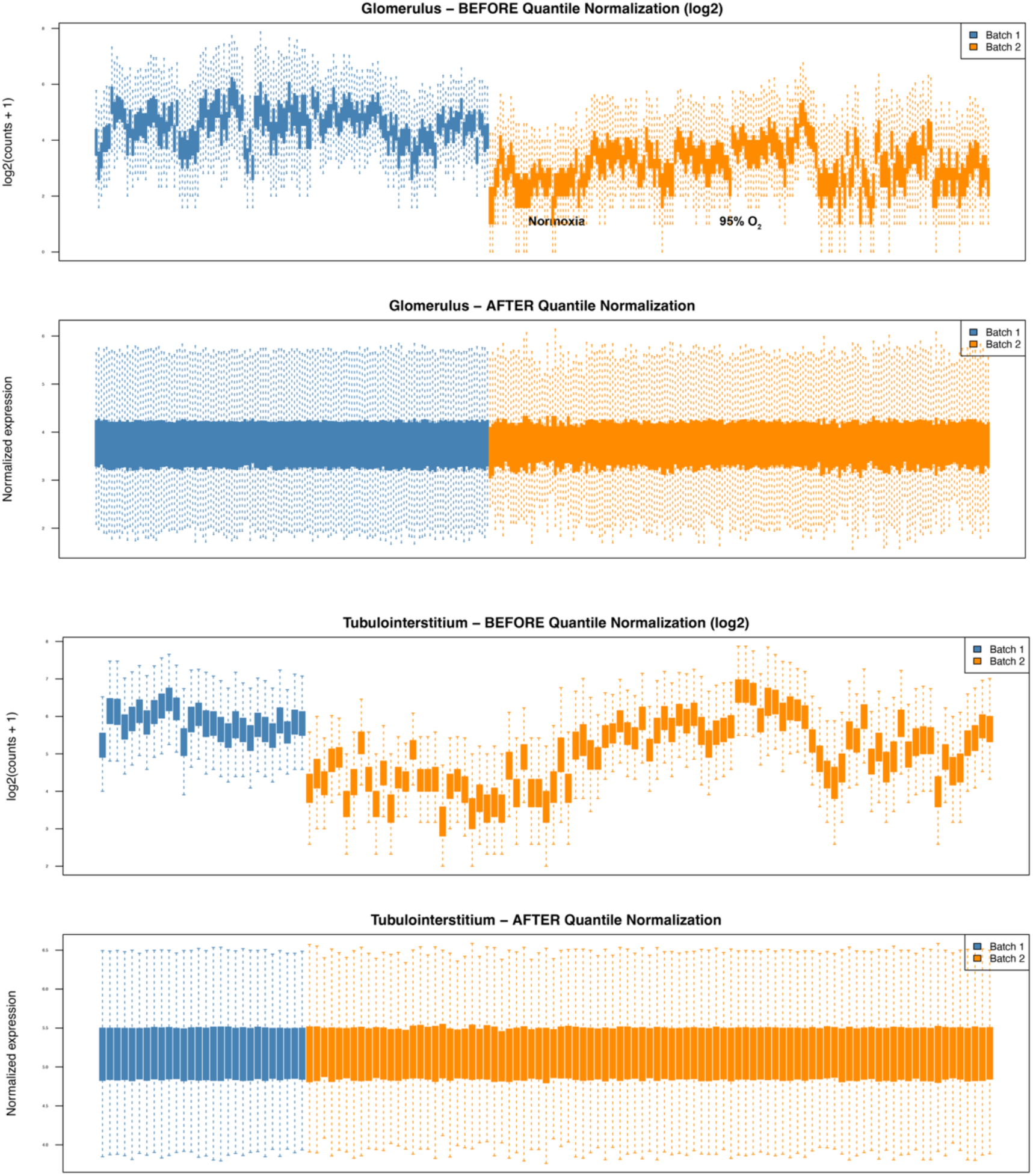
Count distribution in glomerular and tubulointerstitial ROIs before and after quantile normalization.

**Supplemental Figure 3.**
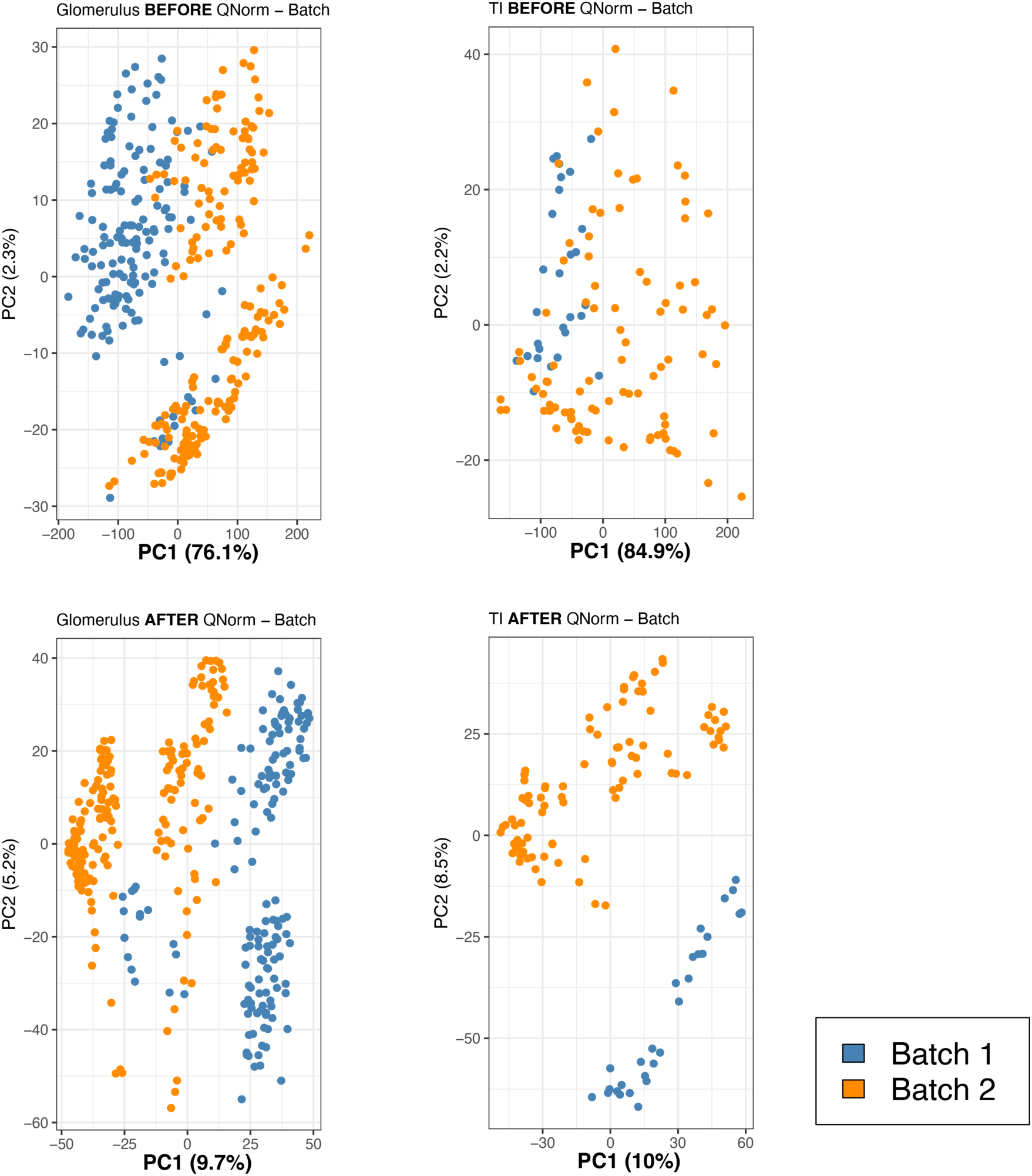
Principal component analysis (PCA) before and after quantile normalization.

**Supplemental Figure 4.**
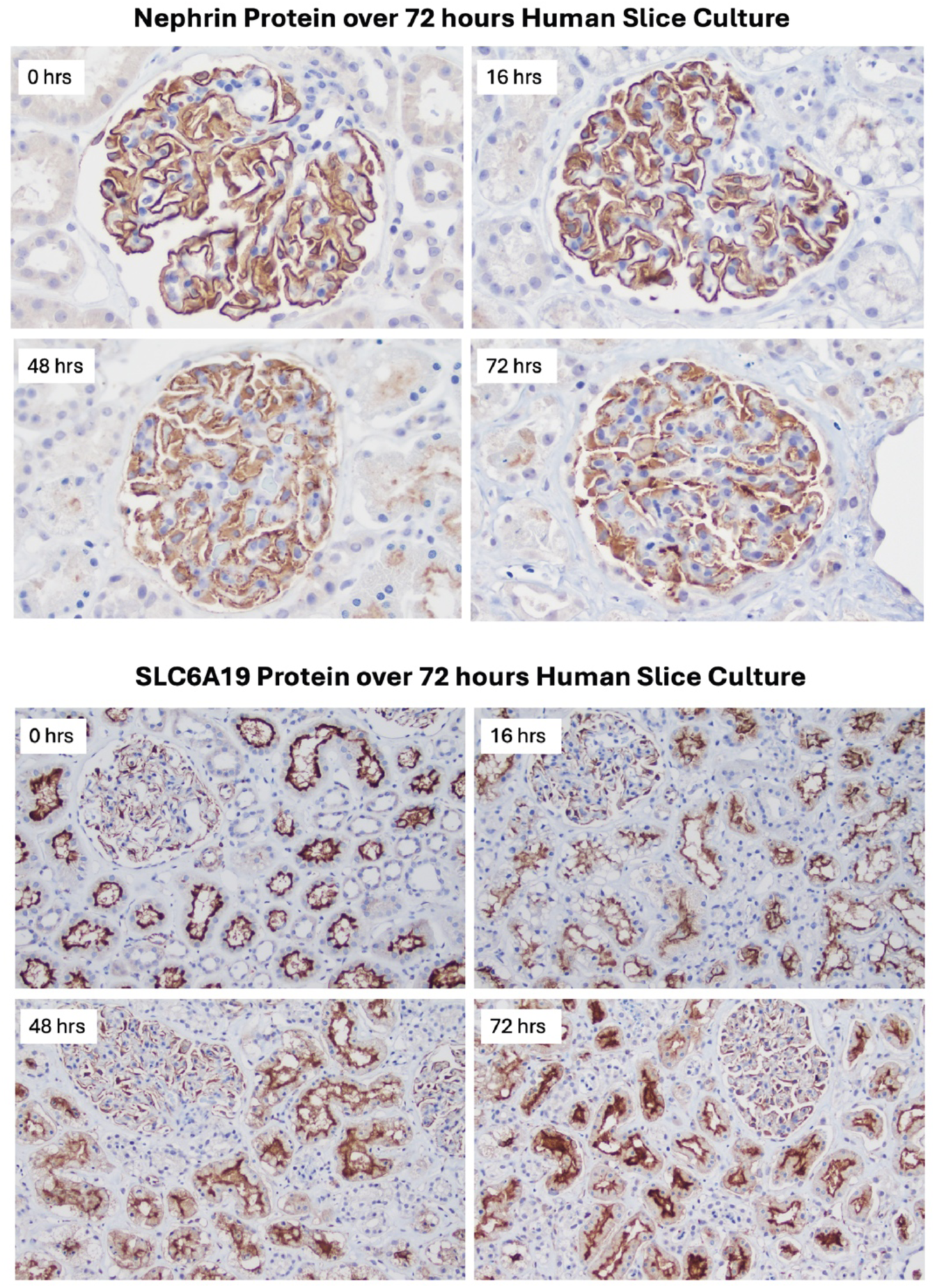
Top, Nephrin (NPHS1) and Bottom, SLC6A19 protein expression over 72 hrs in culture in Medium B-O2.

**Supplemental Figure 5.**
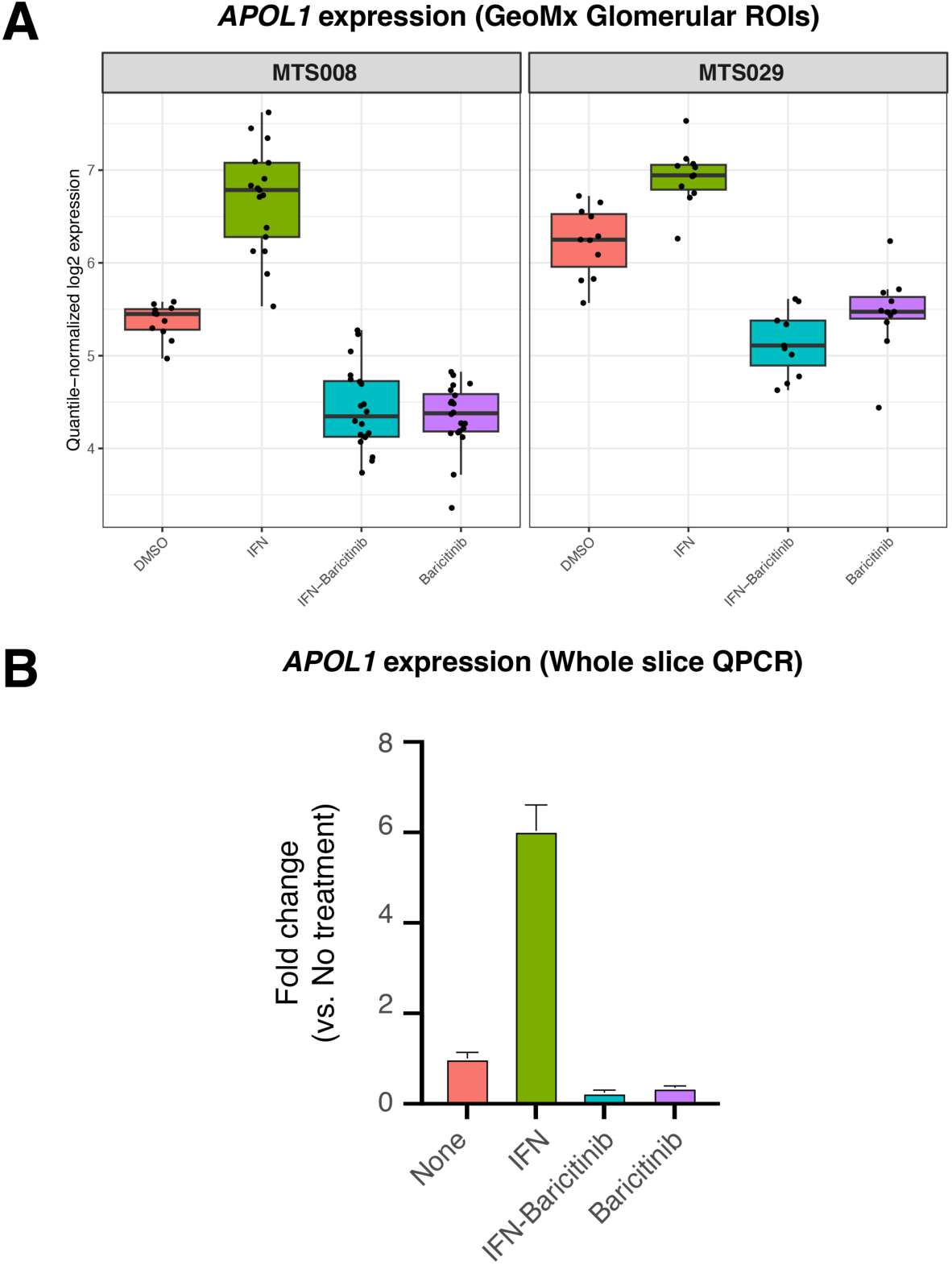
A) Glomerular *APOL1* expression and response to baricitinib after 16 hours in culture. MTS008 was cultured in Medium A, MTS029 was cultured in Medium B-O2. B) Quantitative RT-PCR for *APOL1* normalized to *GAPDH* expression in 3 replicate slices each exposed to the indicated treatments.

**Supplemental Figure 6.**
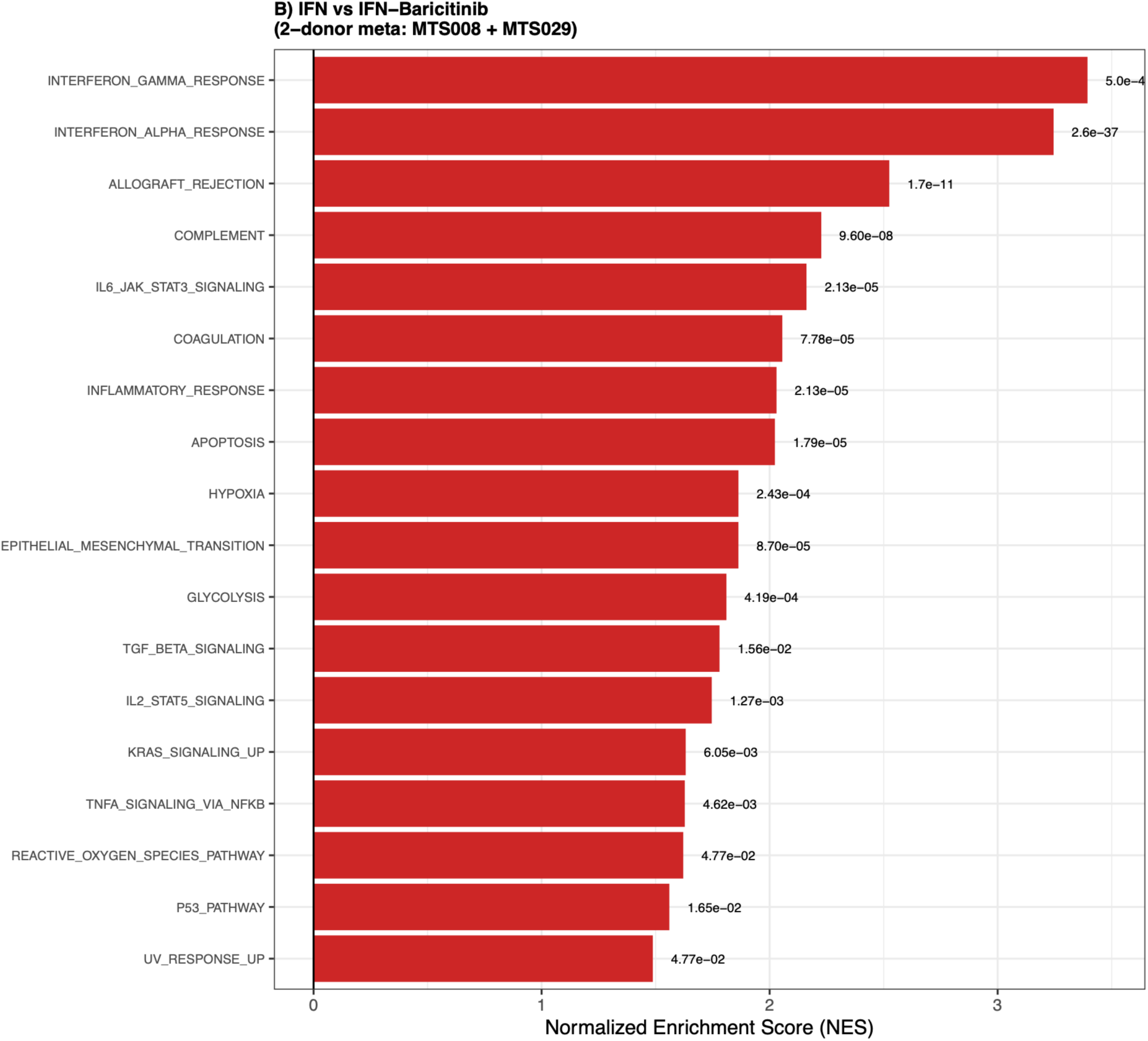
fGSEA of significant enriched pathways in glomeruli following 16 hrs of IFNγ exposure in kidney tissue slices compared to those treated with IFNγ + 1µM baricitinib.

**Supplemental Figure 7.**
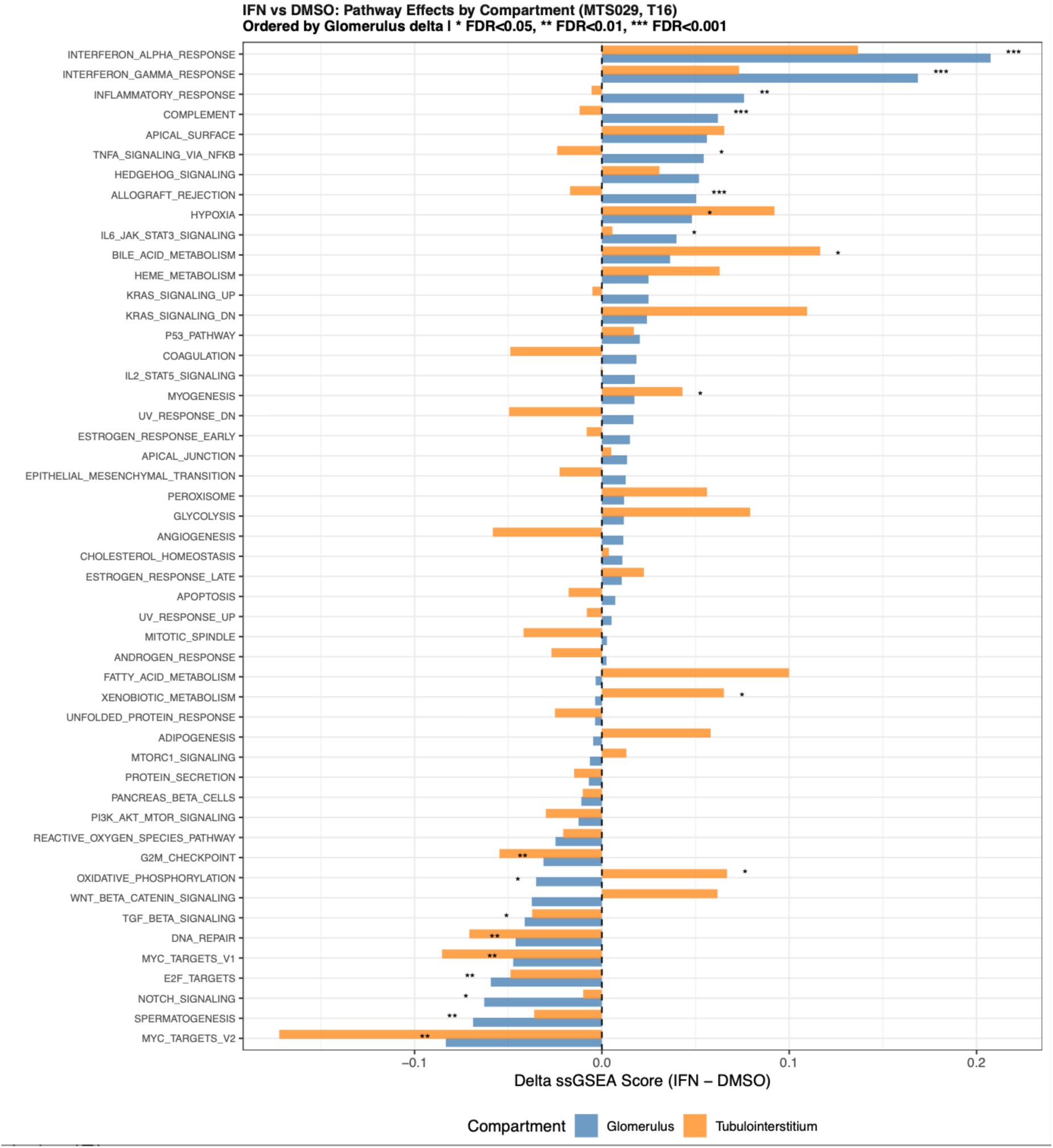
Hallmark pathways impacted by IFNγ treatment in glomerular and tubulointerstitial ROIs.

## SUPPLEMENTAL MOVIES

**Supplemental Movie 1. 3D lightsheet microscopy of a FLARE-stained kidney tissue slice.** Step progression through z-stack of composite image of an entire kidney slice stained using the FLARE approach. Outlines of renal corpuscles containing glomeruli are shaded in red.

**Supplemental Movie 2.** “Fly by” 3D lightsheet microscopy of FLARE-stained human kidney tissue slice highlighting positioning of glomeruli within the slice volume.

## SUPPLEMENTAL TABLES

**Supplemental Table 1. Clinical and pathologic characteristics of kidney tissue donors.** Clinical characteristics and pathologic findings of donors undergoing nephrectomy from whom uninvolved renal cortex was obtained for kidney slice experiments.

**Supplemental Table 2. Primer sequences for quantitative RT-PCR.** Oligonucleotide sequences used for quantitative RT-PCR analyses of kidney slice gene expression.

**Supplemental Table 3. Summary of ROI metadata for GeoMx spatial transcriptomic profiling.** Donor allocation and experimental conditions for two GeoMx Whole Transcriptome Atlas experiments involving eight donors across two batches.

**Supplemental Table 4. Differential gene expression associated with kidney slice culture medium conditions.** Differential expression results for glomerular and tubulointerstitial ROIs comparing freshly prepared tissue (T0) with slices cultured for 16 hours in Medium A or Medium B-O₂ and comparing the two culture conditions. Analyses incorporated donor as a blocking factor.

**Supplemental Table 5. Hallmark pathway enrichment following 16-hour kidney slice culture.** ssGSEA results for Hallmark pathways in glomerular and tubulointerstitial ROIs cultured in Medium A or Medium B-O₂ relative to freshly prepared tissue, with Benjamini-Hochberg-adjusted significance estimates.

**Supplemental Table 6. Top-ranked genes with dynamic expression during 72-hour culture.** Genes with distinct temporal expression patterns underlying the glomerular and tubulointerstitial heatmaps in Figure 5, identified in donor MTS028 cultured in Medium B-O₂ at 0, 16, 48, and 72 hours and ranked by the ANOVA F statistic.

**Supplemental Table 7. Differentially expressed genes in IFNγ-treated glomeruli.** Glomerular differential expression results for IFNγ-treated versus control slices after 16 hours, combined across five donors by fixed-effects inverse-variance-weighted meta-analysis. Cross-donor concordance identifies genes with the same direction of expression change in all five donors.

**Supplemental Table 8. Compartment-specific Hallmark pathway responses to IFNγ.** IFNγ-associated changes in ssGSEA scores in glomerular and tubulointerstitial ROIs, including differences between compartments and FDR-adjusted significance of treatment-by-compartment interactions.

**Supplemental Table 9. Differentially expressed genes in baricitinib-treated glomeruli.** Glomerular differential expression results for baricitinib-treated versus DMSO-treated slices from donors MTS008 and MTS029, combined by fixed-effects inverse-variance-weighted meta-analysis, with assessment of concordant expression changes between donors.

**Supplemental Table 10. Hallmark pathway enrichment following IFNγ stimulation and baricitinib treatment.** Glomerular fGSEA results for IFNγ versus control across five donors, baricitinib versus DMSO across two donors, and IFNγ versus IFNγ plus baricitinib across two donors, including enrichment scores, adjusted P values, and leading-edge genes.

